# Islands of genomic stability in the face of genetically unstable metastatic cancer

**DOI:** 10.1101/2024.01.26.577508

**Authors:** Kirsten Bowland, Jiaying Lai, Alyza Skaist, Yan Zhang, Selina Shiqing K Teh, Nicholas J. Roberts, Elizabeth Thompson, Sarah J. Wheelan, Ralph H. Hruban, Rachel Karchin, Christine A. Iacobuzio-Donahue, James R. Eshleman

**Affiliations:** Department of Pathology, The Sol Goldman Pancreatic Cancer Research Center, The Johns Hopkins University School of Medicine, Baltimore, MD, USA; Institute for Computational Medicine, Johns Hopkins University, Baltimore, Maryland, USA; Department of Oncology, Sidney Kimmel Comprehensive Cancer Center at Johns Hopkins, Baltimore, MD, USA; National Human Genome Research Institute, National Institutes of Health, Bethesda, Maryland, USA; Department of Biomedical Engineering, Johns Hopkins University, Baltimore, Maryland, USA; Human Oncology and Pathogenesis Program, Memorial Sloan Kettering Cancer Center, New York, NY, USA; Department of Pathology, Memorial Sloan Kettering Cancer Center, New York, NY, USA

**Keywords:** CRISPR-Cas9, gene therapy, metastasis, cancer evolution, loss of heterozygosity, rapid autopsy, pancreatic ductal adenocarcinoma

## Abstract

**Introduction:** Metastatic cancer affects millions of people worldwide annually and is the leading cause of cancer-related deaths. Most patients with metastatic disease are not eligible for surgical resection, and current therapeutic regimens have varying success rates, some with 5-year survival rates below 5%. Here we test the hypothesis that metastatic cancer can be genetically targeted by exploiting single base substitution mutations unique to individual cells that occur as part of normal aging prior to transformation. These mutations are targetable because ∼10% of them form novel tumor-specific “NGG” protospacer adjacent motif (PAM) sites targetable by CRISPR-Cas9.

**Methods:** Whole genome sequencing was performed on five rapid autopsy cases of patient-matched primary tumor, normal and metastatic tissue from pancreatic ductal adenocarcinoma decedents. CRISPR-Cas9 PAM targets were determined by bioinformatic tumor-normal subtraction for each patient and verified in metastatic samples by high-depth capture-based sequencing.

**Results:** We found that 90% of PAM targets were maintained between primary carcinomas and metastases overall. We identified rules that predict PAM loss or retention, where PAMs located in heterozygous regions in the primary tumor can be lost in metastases (private LOH), but PAMs occurring in regions of loss of heterozygosity (LOH) in the primary tumor were universally conserved in metastases.

**Conclusions:** Regions of truncal LOH are strongly retained in the presence of genetic instability, and therefore represent genetic vulnerabilities in pancreatic adenocarcinomas. A CRISPR-based gene therapy approach targeting these regions may be a novel way to genetically target metastatic cancer.

## Introduction

Adult stem cells accumulate mutations during the normal aging process(1, 2). Most of these are passenger mutations found in intergenic regions with minimal, if any, impact on cell fitness. Recent studies showed that the rate of accumulation varies based on tissue type but averages approximately 40 mutations per year per cell(3, 4). Over time, this results in a unique set of mutations in each adult stem cell that is clonally propagated to daughter cells during cell division(5, 6). In the event of oncogenic transformation, it follows that the mutational burden in the cancer initiating cell (CIC) should be propagated not only to the growing tumor, but also to metastases that arise from that cancer. These mutations should not be present in the non-neoplastic cells in the same individual. Conversely, since cancer is known to be genetically unstable, it is also possible that cancer-specific mutations that exist in the primary tumor might be lost in some or all metastases.

CRISPR-Cas9 induces site-specific DNA double strand breaks (DSBs) which the cell must repair through endogenous DSB repair pathways(7). As engineered for eukaryotic systems, the endonuclease component, Cas9, is guided to specific DNA target sequences by a single guide RNA (sgRNA)(8, 9).

Cleaving the target DNA by Cas9 is dependent on recognition of a protospacer adjacent motif (PAM) directly downstream of the gRNA target. With SpCas9, the most used CRISPR endonuclease, the PAM recognition sequence is any base followed by two guanine bases, abbreviated “NGG”(10).

Recently, CRISPR-Cas systems have been used for cancer gene therapy. Strategies have included insertion of suicide genes(11), targeting mutations in oncogenic driver genes that contain PAM sites(12, 13), unique sequences in oncoviruses such as E6 and E7(14), and fusion oncogenes(15, 16). We recently reported that somatic single base substitutions (SBS) forming novel PAM sites specific to cancer cells are targetable by CRISPR/Cas9. We discovered that approximately 10% of somatic SBS mutations in cancer cells (that are not present in patient-matched normal cells) produce novel PAM sites. We further demonstrated selective cell killing of tumor cells by exploiting these novel PAMs as targets for CRISPR/Cas9 induced DNA DSBs, finding that targeting 8-12 PAMs is sufficient to reduce PDAC cell populations by about 95%. We showed that patient-matched normal cells were unaffected(17).

Rapid autopsy programs, where tumor tissues are harvested from consented patients within hours of their death, provide an invaluable tool to study cancer heterogeneity and tumor evolution. In this study we used samples from the Gastrointestinal Cancer Rapid Medical Donation Program(18) (PI Dr. Christine A Iacobuzio-Donahue), many of which were established as xenograft tumors or cell lines by our group(19). We sought to identify regions of genomic vulnerability as a precision gene therapy approach. We explored the proportion of novel PAMs present in primary pancreatic tumors that were maintained throughout metastases and the mechanism of loss when PAMs were not maintained. Using deep sequencing of patient-matched primary carcinomas and metastases, we examined evolutionary conservation of PAMs by focusing on the earliest (truncal) mutations. We find that in 5 pancreatic ductal adenocarcinoma (PDAC) cases, an average of ∼90% of truncal PAMs from the primary tumor are maintained throughout each patient’s metastatic lesions and that the primary mechanism when PAMs are lost is loss of heterozygosity (LOH) in metastases. Critically, we find that PAMs occurring in regions of LOH in the primary tumor adjacent to tumor suppressor and oncogenes are 100% maintained in metastases. Counterintuitively, these copy number alterations that contributed to oncogenesis may represent a genetic vulnerability to cancer as essential genes in these conserved regions have been reduced to one copy (with exception of copy-neutral LOH). Building on our previous work, this finding further supports CRISPR/Cas9 as a potential gene therapy for metastatic cancer.

## Results

### Next generation sequencing reveals truncal tumor-specific PAMs in pancreatic cancer

To determine whether CRISPR targeting of tumor-specific somatic PAMs could be used as a potential gene therapy agent against metastatic disease, it is important to understand what proportion of the targets identified in the primary tumor are maintained in each metastasis. The clonal nature of cancer establishes a theoretical foundation for propagation and maintenance of mutations from the CIC. However, cancer is inherently genomically unstable, and such instability might lead to target deletion thereby impacting the efficacy of therapies targeting these mutations.

First, we determined the truncal PAMs for each case, defined here as those PAMs that descended from the original CIC. An experimental and analytical workflow is provided (Figure 1A-B). We performed whole genome sequencing (WGS) on metastatic cell lines derived from five rapid autopsy cases and case-matched normal samples (Table 1). In two of the cases, patients A38 and A32, multiple metastatic cell lines were available for initial whole genome sequencing (WGS) (Figure 1 C-D). In these cases, the union of all PAMs from any of the cell lines sequenced were included for downstream investigation.

**Figure 1:**
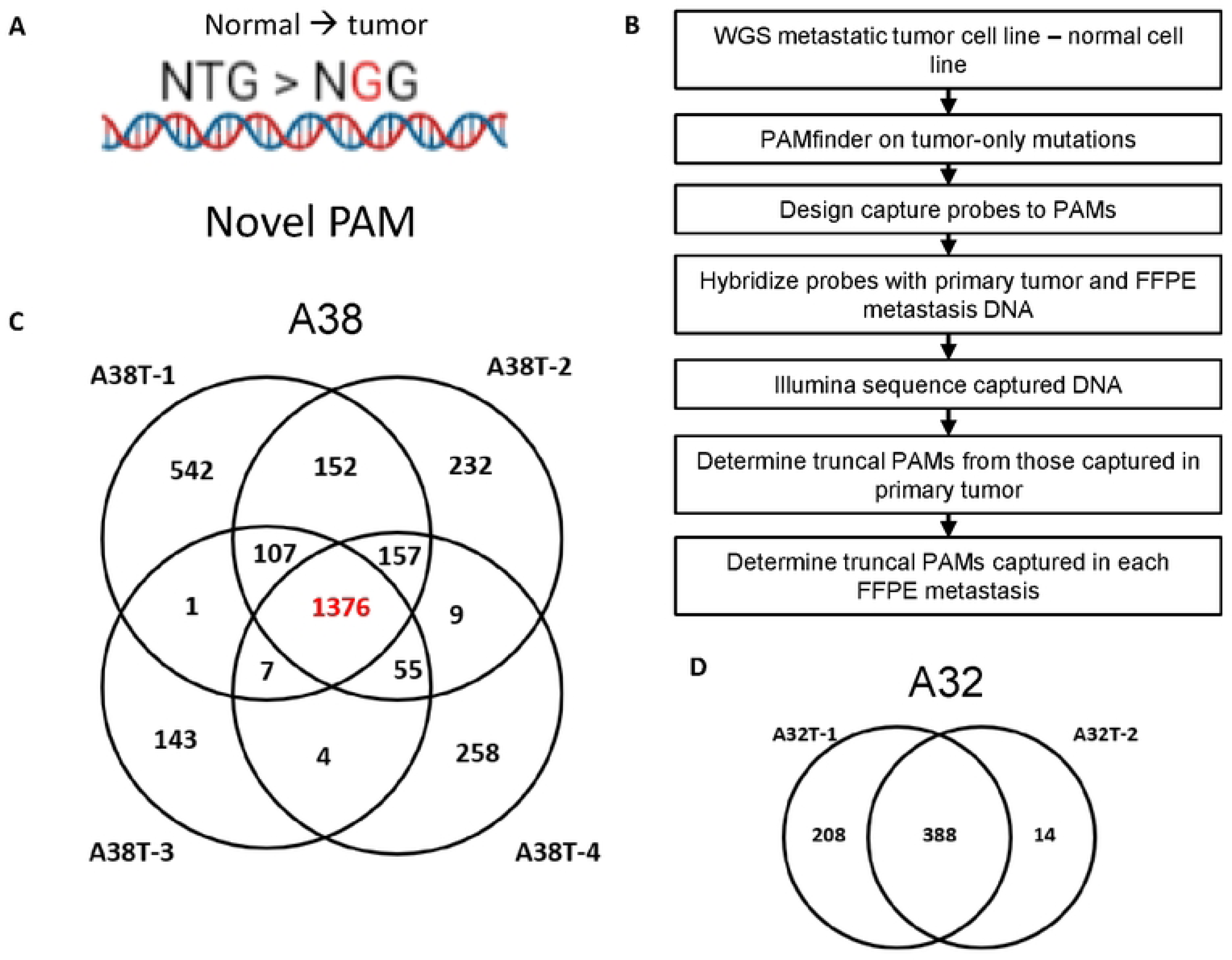
PAM discovery and experimental design. A) Example of novel PAM, where the germline sequence is NTG, and the mutation of T > G forms a novel NGG PAM. **B)** Experimental and analytical workflow. **C)** All PAMs discovered by WGS in the four A38 cell lines and their overlap, **D)** also shown for the two cell lines in case A32.

**Table 1:** Sample summary for whole genome sequencing PAM discovery and PAM validation in metastases.

| **Autopsy Name** | **Source of Cancer Cell Line (WGS Discovery)** | **Source of Normal (WGS Discovery)** | **FFPE Primary and Mets (Validation)** |
| --- | --- | --- | --- |
| **A2** | Liver Metastasis | FFPE liver | Liver mets (5)<br>Mesenteric node |
| **A6** | Liver Metastasis | FFPE spleen | Liver met (2)<br>**Pancreas**<br>Peripancreatic node<br>Omentum<br>Bowel<br>Gall bladder<br>Periaortic mass |
| **A10** | Liver Metastasis | FFPE pancreas | Kidney<br>Testis<br>Periaortic node<br>Cardiac muscle<br>Liver mets (3)<br>**Pancreas (3)**<br>Paraaortic node |
| **A32** | Peritoneal Metastasis<br>Omental Metastasis | EBV lymphs | Diaphragm (2)<br>Omentum<br>**Pancreas** |
| **A38** | Lung metastasis (2)<br>Peritoneal Metastasis<br>Liver Metastasis | EBV lymphs | Unique liver mets (14)<br>**Pancreas (3)**<br>Lung mets (3) |

Tumor-specific PAMs were bioinformatically determined using PAMfinder (Github) after the subtraction of matched tumor and normal variant call files (17). An average of 1236 PAMs (range 379 to 2195) were discovered from each tumor-normal subtraction. PAMs were further analyzed by variant allele fraction (VAF) of 95% or higher to identify a subset of targets in regions of LOH. An average of 217 PAMs (range 18 to 448) were discovered for each subtraction with a VAF ≥ 95% (Table 2). As high quality WGS of primary tumor tissue was unavailable due to formalin fixation, the PAMs captured in any one of the primary samples for each case were considered truncal for this study.

**Table 2:** Tumor minus normal subtractions and the total somatic variants, PAMs, and high VAF PAMs discovered from each subtraction.

| Subtraction | Passed somatic variants | PAMs | PAMs VAF 95%+ |
| --- | --- | --- | --- |
| **A2T-A2N** | 14,644 | 355 | 18 |
| **A6T-A6N** | 28,968 | 1695 | 210 |
| **A10T-A10N** | 18,911 | 664 | 70 |
| **A32T1-A32N** | 16,896 | 614 | 33 |
| **A32T2-A32N** | 9,679 | 379 | 188 |
| **A38T1-A38N** | 21,201 | 2195 | 410 |
| **A38T2-A38N** | 21,210 | 1801 | 312 |
| **A38T3-A38N** | 13,022 | 1645 | 266 |
| **A38T4-A38N** | 16,646 | 1775 | 448 |

### 90% of PAMs from the primary tumor are maintained in metastases

We used DNA capture hybridization probes to investigate the percentage of truncal PAMs that were maintained in multiple metastases for each case (Table 1). As a preliminary analysis, all captured sequences from the primary tumor, normal tissue, and metastases for each case were visualized in Integrated Genome Viewer (IGV)(20) to confirm the presence of the PAM in the tumor samples and absence in the patient-matched normal sample (Figure 2A). As a quality control check for each case, percent truncal for each metastasis was plotted against average read depth for that sample to determine if low cellularity of a particular tissue sample might obscure the presence of a given PAM (Figure S1). For each case, only metastases in the linear portion of these plots were included in the final analysis to compensate for low tumor cellularity or contamination of tumor tissue with surrounding normal cells. For each case, the subset of PAMs present in each tumor sample was plotted as a Venn Diagram to show sample overlap (Figure 2B, S2).

**Figure 2:**
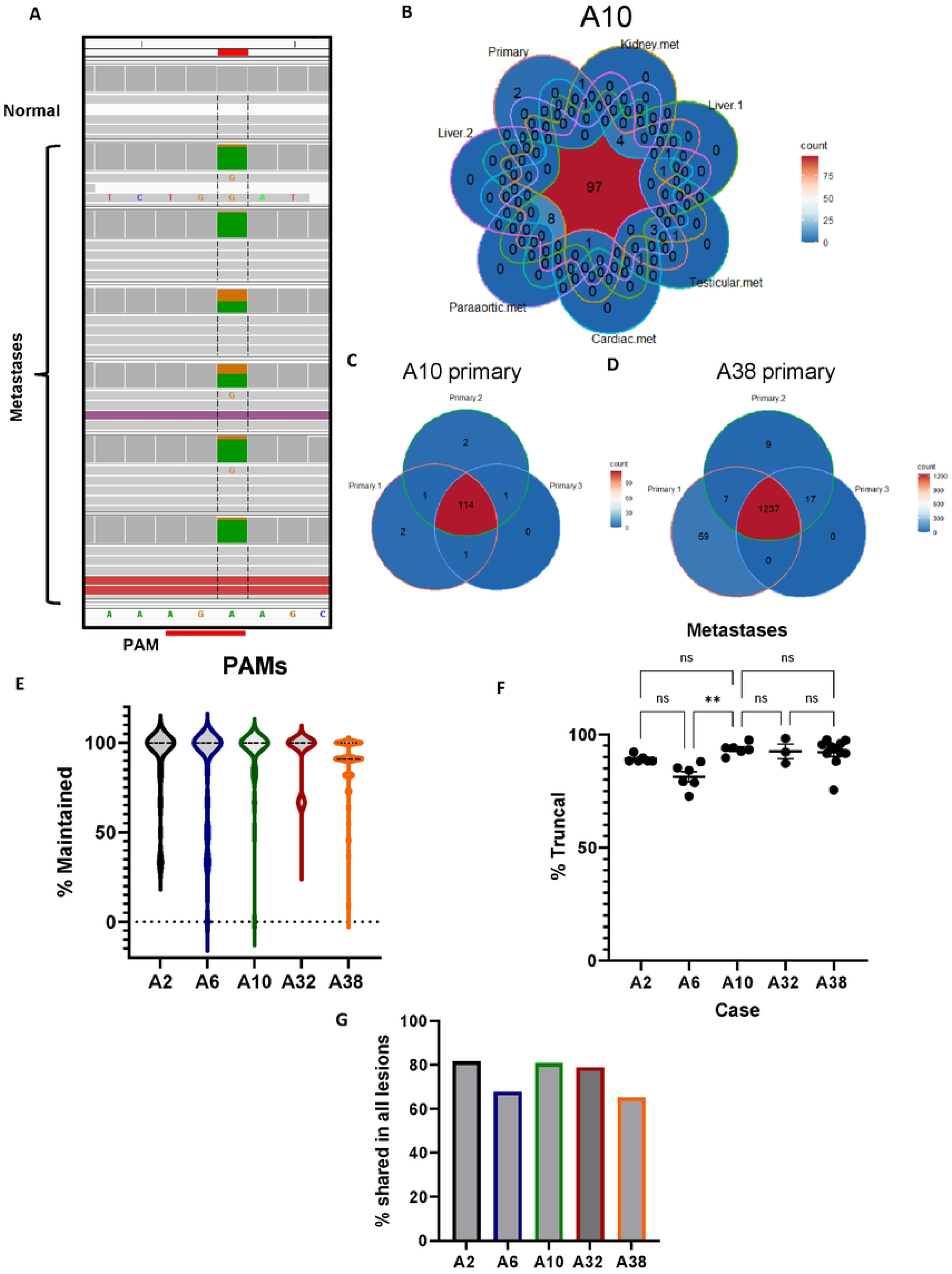
Maintenance of truncal PAMs in metastases. A) Screenshot of capture probe sequencing from IGV at an A>G PAM creating mutation. The PAM is absent in the captured normal tissue but present in all metastatic samples. **B)** Venn diagram of primary and metastatic samples for case A10 showing PAM overlap between all samples. **C)** Overlap of PAMs in primary samples from case A10 and **D)** case A38. **E)** The distribution of metastases in which each PAM in each case is maintained. Each data point is an individual PAM with the mean % truncal and SEM calculated. **F)** The percentage of truncal PAMs that are maintained in each metastasis for each case. Each data point is an individual metastasis. No statistical significance was found between any cases except A6 by one-way ANOVA (p=0.0011). **G)** The percentage of PAMs shared by all lesions in each case.

Two cases, A10 and A38, each had three samples of primary tumor. Using the capture approach, the overlap between those samples was determined, providing insight into heterogeneity within the primary tumor for these cases. A10 primary samples shared 94% of PAMs (Figure 2C). Case A38 shared 90% of PAMs between primary sections but each had distinct PAMs not detected in the other two samples (Figure 2D).

For each case, each metastasis was evaluated for the percentage of truncal PAMs it contained (percent truncal), and each individual PAM was evaluated for how many metastases in which it was present (percent maintained). A2, A6, A10, A32, and A38 had 76, 183, 120, 180, and 1360 truncal PAMs, respectively. These PAMs were maintained in 91.7%, 81.3%, 93.1%, 92.6%, and 88.9% of metastases for each case, respectively (Figure 2E). We also evaluated each metastasis to determine the percentage of truncal PAMs it contained. A2 had 6 metastases and was 89.2% truncal. A6 had 6 metastases and was 81.3% truncal. A10 had 6 metastases and was 93.6% truncal. Case A32, with 3 metastases, was 92.6 % truncal. A38 had 11 metastases and was 92.2% truncal. (Figure 2F). Of note, A2 lacked a sample of primary and so for this case alone, truncal was inferred as mutations present in more than one metastasis. Across all five cases, the average percentage of truncal PAMs maintained in each metastasis was 89.8% and each PAM was present in an average of 89.5% of metastases for each case.

Importantly, for cases A2, A10, A32, and A38, there was no significant difference between the percentage of truncal PAMs in each metastasis by one-way ANOVA. Metastases in case A6 were less truncal than the other four cases (p=0.0016). In addition to examining maintenance of individual PAMs and metastases, the portion of truncal PAMs present in all lesions for each case was also calculated. This includes instances where all but one metastasis shares many PAMs, but a set of PAMs are missing in one lesion. This could be because of focal or regional deletions or LOH. Examining each case this way, we find that for A2, A6, A10, A32, and A38, 81.6%, 67.8%, 80.8%, 78.9% and 65.1% of PAMs were shared between all lesions studied, respectively. Across all five cases, an average of 74.8% of PAMs discovered in the primary tumor were present in every lesion studied (Figure 2G).

### PAMs are 100% maintained in regions of truncal LOH proximal to tumor suppressor and oncogenes

A deeper analysis of copy number (CN) and LOH was performed on cases A32 and A38, as WGS of two and four tumor cell lines, respectively, and matched normal cell lines were available.

Zygosity and copy number directly affected the PAM VAF in sequencing data as diagrammed (Figure 3A). We then mapped chromosomal location of PAMs together with their VAF and copy number for each tumor cell line and found that truncal PAMs were more likely to have a VAF of 100% and be in haploid regions (Figure 3B-C, S3-S4).

**Figure 3:**
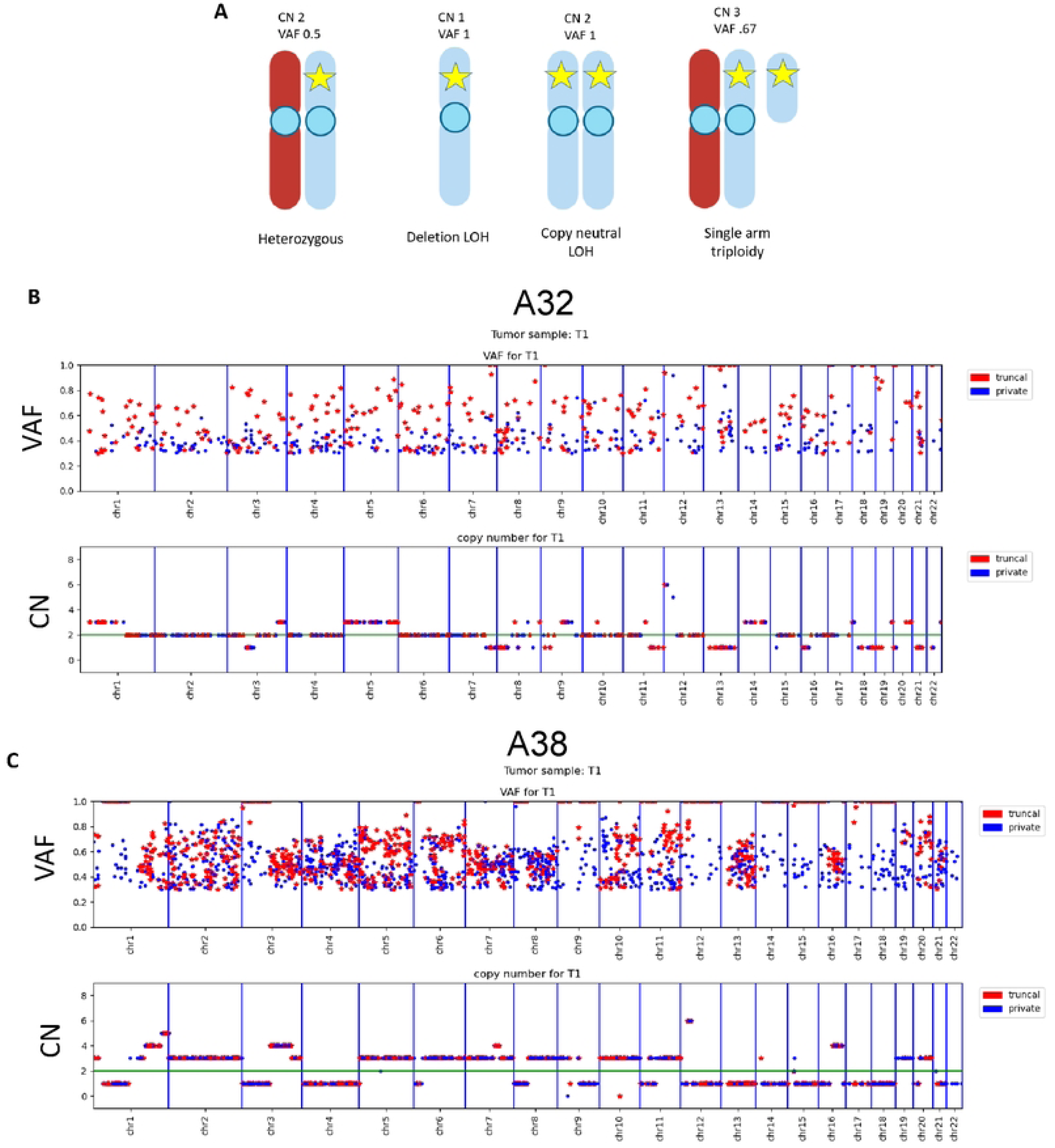
Truncal PAMs have higher VAFs than private PAMs. A) Diagram depicting relationship between variant allele fraction (VAF), ploidy, and copy number (CN) when interpreting next generation sequencing data. WGS from cell lines show truncal PAMs with high VAF frequently occur more frequently in haploid regions of the genome. Shown is **B)** A32-T1, representative for case A32, and **C)** A38 cell line T1, representative for case A38.

To investigate the relationship between VAF, copy number, and PAM maintenance further, we cross-referenced the list of PAMs found in all three captured samples of the A38 primary tumor with the WGS cell line data. First, we quantified the number of truncal PAMs on each chromosome by VAF and copy number (Figure S5-S6). We found that on chromosomes where all PAMs were present at a VAF of 100%, all PAMs were present in all four tumor cell lines (Figure 4A-B). On chromosomes which were haploid in one or two cell lines but not all, indicating that the copy loss occurred later in the disease, we found that PAMs were partially lost in the haploid regions. PAMs were 100% maintained in the regions that retained heterozygosity (Figure 4C-D). Furthermore, PAMs were 100% maintained on chromosomes that were diploid or greater and heterozygous in all four cell lines (Figure 4E-F).

**Figure 4:**
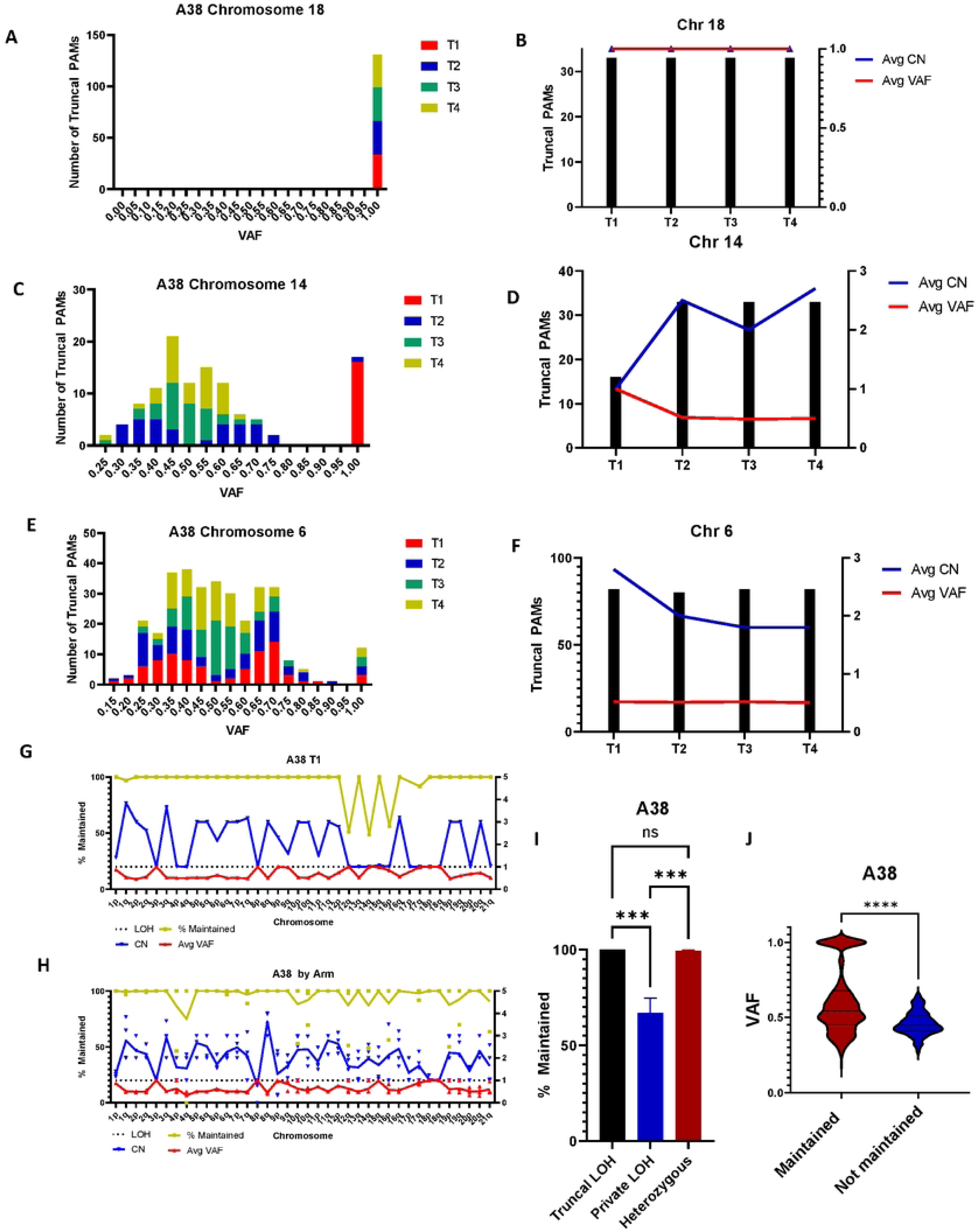
PAMs are maintained in regions of truncal LOH and partially lost in regions of private LOH. All data shown is from case A38. **A)** All truncal PAMs on chromosome 18 in all four cell lines have a VAF of 1 and **B)** all truncal PAMs discovered are present in all four cell lines with a copy number of 1 (haploid). **C)** A38 cell lines T2-T4 are at least diploid on chromosome 14 while T1 is haploid based on VAF. **D)** Truncal PAMs are 50% lost in T1, corresponding to a VAF of 1 and a CN of 1. PAMs are 100% maintained in T2-T4 where the VAF remains 0.5, indicative of heterozygosity. **E)** Chromosome 6 is at least diploid in all cell lines. Notably, T1 and T2 are triploid as indicated by VAF peaks at 0.35 and 0.7. **F)** All truncal PAMs are maintained in all four cell lines as heterozygosity is maintained. **G)** Percent PAM maintenance, average CN, and average VAF plotted by chromosome arm for A38 cell line T1. PAM loss occurs in haploid regions, where the VAF and CN are both 1. **H)** PAM maintenance, average VAF, and average CN across chromosome arms aggregated for all four A38 cell lines. Notably, areas of LOH common to all cell lines (truncal LOH) present 100% PAM maintenance, while PAM loss is limited to regions of private LOH. Regions of truncal LOH can be identified where CN=1 in all samples and/or VAF= 1 in all samples. **I)** Summary data of truncal PAMs in case A38 WGS data. Truncal PAMs are 100% maintained in regions of truncal LOH, 67% maintained in regions of private LOH, and 99% maintained in heterozygous regions. Pairwise comparisons by Kruskal-Wallis test. **J)** Plot of truncal PAM VAF comparing PAMs maintained in all four cell lines and truncal PAMs lost in at least 1 cell line. Truncal PAMs that are maintained have a significantly higher mean VAF (0.6029) than those susceptible to loss (0.4598) (Mann-Whitney Rank Sum, p=2.267x10^-16^). Additionally, maintained PAMs are more likely to have a VAF of 1 indicative of their occurrence in regions of LOH. In contrast, PAMs that were lost had a maximum VAF of 0.6483. n=1080 for maintained PAMs, n=146 for lost PAMs. All error presented in SEM.

We then examined the genome of each metastasis, calculating the average PAM VAF, CN, and the percentage of maintained PAMs for each chromosome arm. Strikingly, PAMs were overall maintained, or lost at a rate of about 50% per copy, i.e., 50% lost in haploid regions and 33% lost in polyploid regions (Figure 4G). Aggregating data from all four A38 cell lines, we found that PAM loss was limited to regions of private LOH, where individual samples had undergone LOH, but others had not (Figure 4H, S7). Further examining patterns of PAM maintenance and loss, we found that PAMs were 100% maintained in areas of truncal LOH but lost at a rate of 30% on chromosome arms that contained private LOH. Likewise, in A32, PAMs were 100% maintained in regions of truncal LOH, and were only lost in regions of private LOH or regional deletions (Figure S8). In A38, PAMs are maintained at a rate of 99% where heterozygosity is maintained, indicating private LOH as the primary mechanism of PAM loss (Figure 4I, S9A). Further supporting this hypothesis, we found that truncal PAMs that were 100% maintained had a higher average VAF and are more likely to be in regions of haploid LOH than truncal PAMs that had been lost in at least one metastasis (Figure 4J). A similar trend was seen in case A32, although statistical significance was limited by a small number – only 3—of lost truncal PAMs (Figure S9B).

Finally, when the percent maintained was plotted as a function of VAF, we saw that truncal PAMs maintained in all four A38 cell lines have a broad VAF distribution. Truncal PAMs lost in at least one cell line, however, were 100% maintained at a VAF of 50% (regions of retained heterozygosity), or 50% lost at a VAF of 100% (regions of LOH), supporting the hypothesis that LOH was responsible for loss of individual PAMs (Figure S9C).

## Discussion

### Haploid conserved islands of essential genes represent a therapeutic vulnerability

In both cases for which we possessed cell lines from multiple lesions, A32 and A38, we observed haploid regions that were consistently maintained between the primary tumors and metastases. Unlike other areas of private LOH representing ongoing genomic instability, these regions were haploid in the primary tumor and 100% conserved in metastases. These regions of truncal LOH frequently occurred around tumor suppressor and oncogenes commonly mutated in PDAC and likely contributed to transformation in the cancer initiating cell. Counterintuitively, these regions may represent a therapeutic vulnerability, as many genes shown to be essential in PDAC are also located in these haploid islands(21). LOH islands surrounding mutated oncogenes like KRAS should be under selective pressure if oncogenic addiction is present to the driver gene.

In case A38, the areas of truncal LOH included chromosome arms 1p, 3p, 8p, 9p, 15q, and the entire chromosome 18. Chromosome 8p was largely deleted in cell line A38 T2, but the regions containing truncal PAMs were conserved. These chromosome arms contain tumor suppressor genes frequently deleted or mutated in pancreatic cancer, such as CDKN2A and *SMAD4*. In A32, regions of truncal LOH were present on chromosomes 12p and 20p. Chromosome 12p harbors activating KRAS mutations in the majority of PDACs, including A32. It is possible that targeting PAMs in regions of haploid LOH proximal to mutated tumor suppressor genes would ensure that the targeted PAMs are maintained in all metastases, and further increase the toxicity of the incurred DSBs as those regions lack a template for homology directed repair of the DNA lesion.

### CRISPR-Cas9 guided tumor-specific DSBs as a cancer gene therapy

Our previous work and that of others showed that a limited number of double strand breaks is catastrophic for cancer cells(22–24). Mechanistically, our group has found that multiple simultaneous DSBs trigger the formation of cytogenetic lesions such as radial, dicentric, and ring chromosomes. These lesions lead to catastrophic genomic instability and cell death after a period of 7 to 21 days(25).

Our group recently performed proof of principle work to demonstrate CRISPR-Cas9 as a targeted cell killing tool against cancer(17). Here we find that 90% of tumor targets can be found in the primary tumor and metastases, demonstrating that our approach should be applicable for treating metastatic cancer. *In vivo* studies with mice xenografted with patient derived cancer cells and patient derived organoids would be helpful to demonstrate cell killing of multiple tumors with the same set of targets. Also, *in vivo* delivery of CRISPR agents is still under investigation by many researchers(26) and much optimization is needed(27). However, one could envision an approach with lipid nanoparticle encapsulation of Cas9 mRNA and sgRNAs, similar to the COVID-19 vaccine developed by Moderna and Pfizer(28). Moreover, as researchers dedicated to drug delivery optimize *in vivo* approaches for a myriad of therapeutic purposes, it is critical that basic science researchers develop therapies in parallel so that the two fields will meld in the future to the benefit of patients.

### Implications of PAM maintenance on patient sampling

Determining the clonal PAMs in each metastasis should indicate how many biopsies must be taken from a patient to ensure precision targeting of a patient’s entire disease. As we show here that 90% of truncal PAMs were indeed present in metastases, it should not be necessary to extensively biopsy a patient to narrow down the set of targetable mutations to generate enough DSBs in each cancer cell to achieve cell death. Additionally, we found that PAMs were maintained at a rate of 100% in areas of truncal LOH. In these cases, the PAMs may be selected for in metastases, as loss of the PAM would mean loss of all genetic information in that region. Identifying PAMs in regions of LOH in the primary tumor should then bias the analysis towards PAMs that will be maintained in all metastases. This conclusion is supported by a recent study which found that mutations in haploid or amplified regions of the genome constituted a persistent mutation burden in cancer which was more likely to be maintained(29).

Pancreatic cancer metastases can be polyclonal in origin(30), so sampling multiple sections of the primary tumor might elucidate much of this primary tumor heterogeneity passed on to metastases. Computational modeling supports this conclusion. A recent study using spatial modeling of intratumoral heterogeneity predicted that 3 -5 biopsies of a primary tumor were sufficient to identify clonal mutations, depending on tumor heterogeneity and clonal complexity(31). Similarly, sequencing of homogenized tumor samples from surgical pathology “waste” was shown to reveal heterogenous mutations present in primary tumors(32). One could envision our approach as an adjuvant treatment targeting residual disease after primary tumor resection – in such a scenario, multiple sequencing samples from the primary tumor should be sufficient to reveal the truncal PAMs. In cases where a patient is diagnosed after metastatic spread, which is frequently the case, or as recurrent disease after resection, a small number of biopsies should be sufficient to distinguish truncal PAMs from other subclonal PAMs and allow targeting of all metastatic sites.

### Maintenance of truncal PAMs in metastases

Most studies focused on the appearance and propagation of driver mutations in tumor evolution. Previous work with case A38, published as sample Pa08, revealed a stepwise progression of driver mutations occurring subclonally in the primary tumor, allowing for clocking of initial tumorigenesis and metastatic spread(33). A recent study of clonality in melanoma metastases revealed multiple metastatic lineages with distinct mutational profiles(34). Here, however, we focused on passenger mutations, largely intergenic, which were subject to less selective pressure than drivers.

Frequent events such as LOH, chromosomal deletions, and structural variants could also remove or affect the copy number of truncal PAM sites. In the context of CRISPR-based gene therapy, these alterations could impact how many DNA DSBs would be generated per target or the choice of DNA repair pathway employed after the break(35), affecting downstream cytotoxicity. To cause the highest cytotoxicity with the smallest number of DSBs to the most cancer cells, it is crucial to understand the genomic landscape of novel PAMs throughout a patient’s disease.

For the purposes of this study, we defined “truncal” as those mutations present in the primary tumor. Truncal has been previously defined as “ancestral mutations in the trunk of the phylogenetic tree that are shared by all clones” (36). While it is impossible to sequence the CIC, clonal mutations can be inferred from multiple samples of a primary tumor. Here we included PAMs discovered in any sample of primary tumor. Targeted deep sequencing of sonicated DNA has been shown to introduce spurious SBS mutations in sequencing data(37) and somatic PAMs are frequently located in repetitive areas of the genome leading to poor probe specificity and low sequencing coverage of those regions. Multiple samples of primary tumor were available for two cases (A10 and A38). To maximize the number of PAMs considered for this study and to eliminate the chance that the PAMs are present throughout a primary tumor but not detected because of technical artifacts, all PAMs discovered in any sample of the primary tumor were included for downstream analysis of clonality.

There are some limitations to this study. Cell lines were used to generate whole genome sequencing to identify somatic PAMs, and these are not available in all cases of cancer. Ideally, one would determine this information by performing whole genome sequencing (WGS) directly on primary tumor samples, however in some cancers like pancreatic cancer, such analysis can be compromised by low primary neoplastic cellularity. Analyses can be further complicated by sample age, oxidation, and crosslinking in formalin fixed paraffin embedded (FFPE) blocks. To begin with the largest pool of PAMs possible, we sequenced high quality DNA from metastatic cell lines for each patient and designed capture probes complementary to the combination of all PAMs found in each case. Naturally such a method included many private mutations for each metastasis, including those that arose during passage in cell culture. We were able to narrow down the broad pool of mutations for each case to those specific to the primary tumor by hybridization with FFPE primary samples. It is possible that some truncal PAMs present in other subclones of the primary tumor not evolutionarily upstream of the WGS metastatic cell lines were excluded with this approach and that starting analysis with PAMs already propagated to metastases biased the analysis. However, CIC mutations should be present in every subclone of the primary tumor prior to metastasis, and so isolating the mutations present in multiple metastases and the primary tumor helped limit this potential bias.

As LOH is an event unique to cancer, investigators have attempted to exploit this signature using various methods for some time(38). Recently, Kinzler et al. explored a CAR-T immunotherapy approach targeting LOH at HLA alleles in cancer(39). As PAMs are preferentially maintained in haploid regions of LOH, our approach allows for prediction of targets that could be efficacious in treating metastatic disease. Identification of conserved islands of LOH surrounding known tumor suppressor and oncogenes has implications beyond our study and warrant further investigation. Our approach may allow for genetically targeting LOH in cancer for a range of therapeutic applications.

## Materials and Methods

### WGS PAM Discovery

DNA was extracted from cell pellets or frozen tissue with a Qiagen UCP Micro DNA Kit (Ref 56204) per the manufacturer’s instructions. Matched tumor and normal samples were verified by STR profiling. WGS was performed on cancer samples to a target depth of 60x coverage, and 30x coverage for normal tissue. Sentieon-genomics-202010.02 bwa v0.7.17 (mem) was used for running the alignments against hg19 genome using default parameters. Sentieon-genomics-202010.02’s THhaplotyper was used to call somatic variants between the tumor-normal pairs. CNVkit v0.9.4 (batch) was used for copy number variation detection using “wgs” as a method parameter. The resulting integer copy numbers in the call.cns files from teh CNVkit outputs were used for downstream analysis. Sentieon-genomics- 202010.02’s Haplotyper was used to call germline variants in normal samples with default parameters. The variants were filtered by the mapping quality ≥20, depth ≥10 and an allele frequency ≥0.05.

PAMfinder (previously described) was then used to call somatic PAMs specific to tumor samples. PAMfinder is available for download at https://github.com/selinateh/PAMfinder.

### FFPE Tissue Macrodissection

5 µM H&E slides were cut from each FFPE rapid autopsy blocks and reviewed by an anatomical GI pathologist (ET) with marking of tumor and normal regions prior to macrodissection. 10-µM sections were cut adjacent to the H&E section and were not stained. Sections were deparaffinized with xylene and ethanol. The PinPoint slide DNA isolation system (Zymo Research, USA) was used according to manufacturer’s directions to dissect and digest desired areas of tissue. Samples were digested with proteinase K overnight and DNA was extracted the next day by QIAamp DNA Mini Kit (ref 51306) (Qiagen, USA) with washes performed twice. DNA was quantified by Qubit HS Assay (ThermoFisher Scientific, USA).

### Capture Probe Design

xGen LockDown Hybridization Probes (IDT, USA) were custom designed. Probe pools consisted of 120bp biotinylated oligonucleotides with 2x tiling over the desired areas, where possible. Probes were designed to straddle novel PAMs found from PAMfinder script as described above. To prevent nonspecific binding, probes were not included for loci with specificity scores greater than 100 as determined by the manufacturer’s algorithm (range 1-500, >50 considered risky). Samples were broken into two pools, with A2 and A32 combined in Pool 1 (3379 probes) and A38, A10, and A6 in Pool 2 (9239 probes).

### Library Preparation

Macrodissected DNA samples were sheared on a Covaris S2 Focus Ultrasonicator (Covaris, USA) to a target length of 250-350 bases. Libraries were prepared by xGen cfDNA & FFPE DNA Library Prep kit (IDT, USA) per manufacturer’s instructions. Libraries were amplified with KAPA HiFi HotStart Ready Mix (Roche, USA) and bead purified by AMPure XP beads (Beckman Coulter, USA). DNA was quantified by Qubit HS Assay or D1000 High Sensitivity Screen Tape (Agilent, USA).

### Hybridization

Libraries were multiplexed with equal mass ratio in 4, 5, or 8-plexes. Combined libraries were dried down with universal blockers and human cot DNA in a Savant SpeedVac (ThermoFisher Scientific) vacuum concentrator. Dried down samples were hybridized overnight with custom panels of xGen LockDown Probes (IDT, USA). Washes and bead captures were performed the following day with xGen Hybridization reagents per the manufacturer’s instructions. Captured libraries were amplified with KAPA HiFi HotStart Ready Mix (Roche, USA) and purified with AMPure XP beads. Final capture libraries were quantified on a Bioanalyzer (Agilent, USA) prior to sequencing.

### Capture Library Sequencing

Capture libraries were sequenced with 2x250 paired end reads on an Illumina MiSeq to a target depth of 100x coverage. SAMTOOLS (RRID: SCR_002105) v1.10 and GATK v3.6.0 were used to determine coverage at different levels of partitioning and aggregation.

### Analysis of Capture Data

Aligned reads were visualized in Integrated Genome Viewer (IGV, version 2.15.4, Broad Institute)(20) with single base substitutions called at 5% VAF cutoff, just above the threshold for background artifactual noise. Metastases were visualized concurrently with captured normal and primary tumor sequencing for each case to determine the presence or absence of each PAM. All PAMs present in the primary tumor capture but absent in captured normal were considered truncal. “Percent truncal” for each metastasis was calculated as the number of truncal PAMs present in that sample divided by the number of truncal PAMs for the case. “Percent maintained” was calculated for each PAM by dividing the number of metastases in which the PAM was present by the total number of metastases for that case.

### Statistical analysis

All statistical analysis was performed with GraphPad Prism (RRID: SCR_002798) software 8.3.0. Nonparametric comparisons of three or more groups were determined with Kruskal-Wallis test and nonparametric comparisons of two groups were performed with Mann-Whitney Rank Sum test. Results were considered significant at p < 0.05. Error is displayed as SEM throughout the study.

### Sex as a biological variable

For the analysis of PAM maintenance in metastasis, 80% of patients were male and 20% were female. Patients A2, 6, 10, and 38 were male. Patient A32 was female. For the WGS analysis of LOH islands and mechanism of PAM loss, 2 total cases were analyzed, one of which was male (A38) and one of which was female (A32).

## Acknowledgements

The authors would like to thank the patients and their families who consented to rapid autopsy and made this study possible. We would also like to thank Jennifer Meyers, Drs. Kornel Schuebel, and Srinivasan Yegnasubramanian, Laura Wood, Suping Chen, Richard Burkhart, Chien-Fu Hung, Angelo DeMarzo, and Ming-Tseh Lin for helpful suggestions; Arnique Coutain, Julia Mychak, Lisa Haley for technical assistance; Kailee Calder, Fidel Cai, Jonathan Chen, and Kindness Nwaukwa for assistance with data analysis.

## Data availability

The datasets generated and/or analyzed during the current study are available from the corresponding author on reasonable request.

## Ethical approval

Experiments were performed under IRB approval.

## Additional information

- **Financial support:** The Stringer foundation (JRE), Susan Wojcicki and Dennis Troper (JRE), The Sol Goldman Pancreatic Cancer Research Center (JRE), National Institutes of Health grant P50CA62924 (Dr. Alison P. Klein), National Institutes of Health grant P30CA006973 (Dr. William G. Nelson), and the Mary M. Graf, Linda C. Talecki, Casimir H. Zgonina, Elaine Crispen Sawyer, Eve Stancik, George Rubis, Professor J. Mayo Greenberg and Dr. Samuel L. Slovin, James S. McFarland, Hilda B. Yost, Mary Lou Wootton, Dick Knox/Cliff Minor, John J. Lussier, Rawlings Family, Edward Goldsmith and Elaine T. Koehler Cancer Research Foundations.
- **Corresponding author:**James R. Eshleman, MD, PhD; Department of Pathology and Oncology, 1550 Orleans Street, Suite 344, Baltimore, MD, 21231; 410-955-3511 (phone), 410-614-4734 (fax).
- **Author contributions:**

`o` Kirsten Bowland: conceptualization, investigation, formal analysis, initial manuscript preparation
`o` Jiaying Lai: investigation, formal analysis
`o` Alyza Skaist: formal analysis
`o` Yan Zhang: formal analysis
`o` Selina Shiqing K Teh: conceptualization, manuscript editing
`o` Nicholas J Roberts: conceptualization
`o` Elizabeth Thompson: formal analysis
`o` Sarah J. Wheelan: formal analysis
`o` Ralph Hruban: conceptualization, manuscript editing
`o` Rachel Karchin: investigation, formal analysis
`o` Christine A Iacobuzio-Donahue: conceptualization, sample procurement
`o` James R. Eshleman: conceptualization, oversight, funding procurement, manuscript editing

## Conflict of interest disclosures

Drs. Bowland, Teh, Roberts, and Eshleman, and Johns Hopkins University filed a provisional patent with the USPTO.

---

**Supplemental Figure 1.**
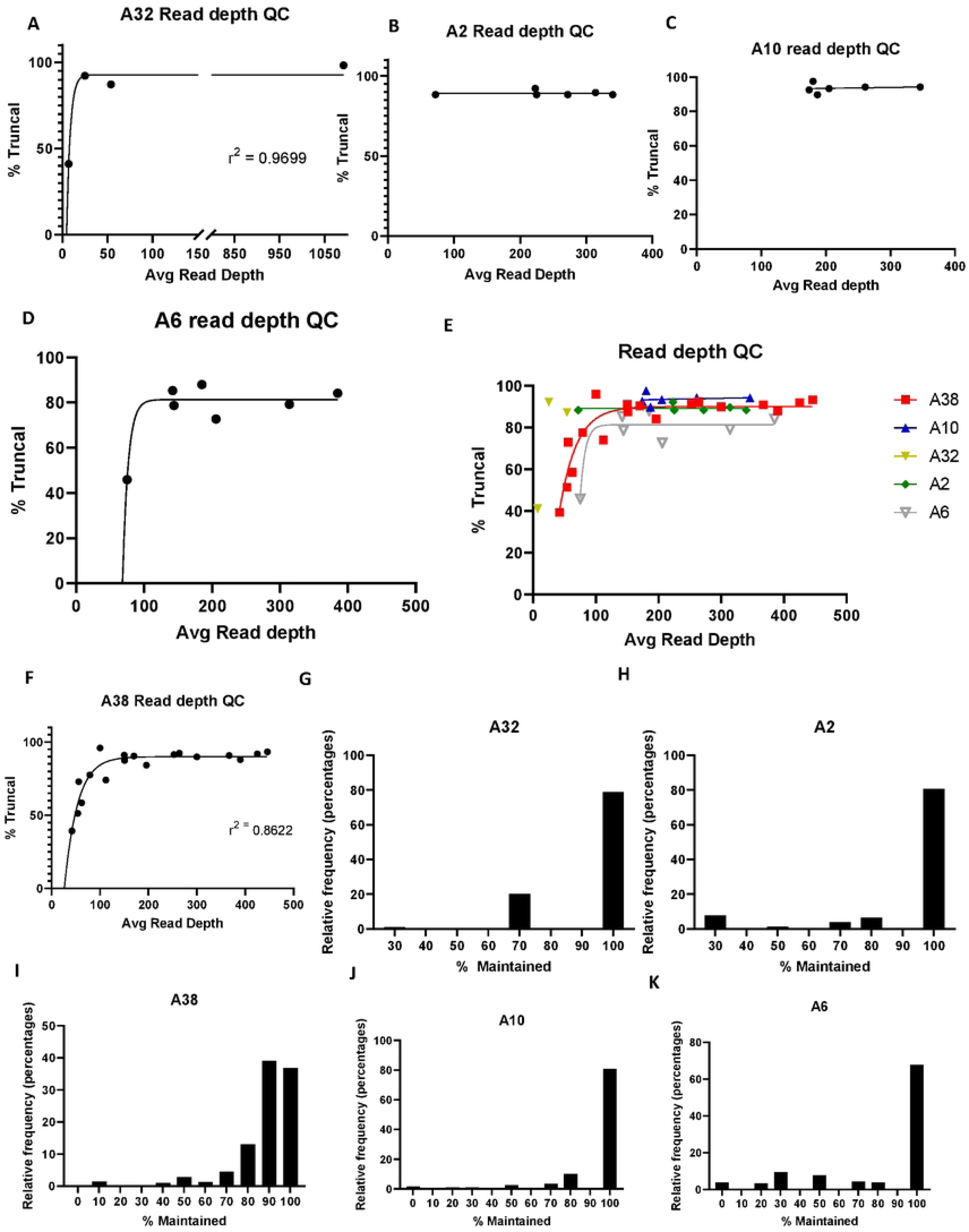
Quality control analysis plotting average read depth for each sample against the calculated percent truncal for that sample, broken down by each case, and then shown overall for all cases. Histograms showing maintenance of PAMs for each case are also shown.

**Supplemental Figure 2.**
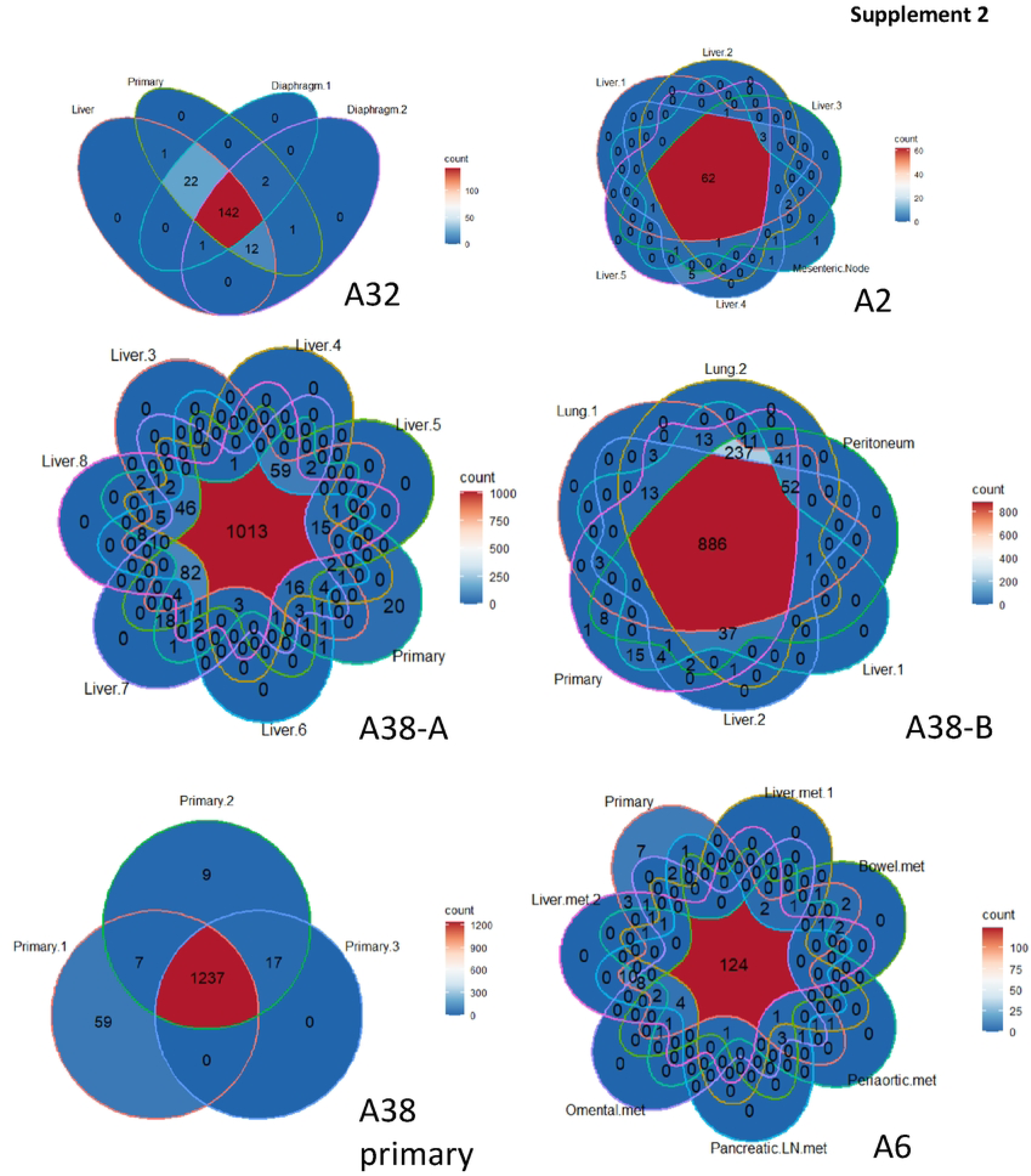
Venn diagrams of PAM overlap for each case. A38 is broken into two Venn diagrams as a maximum of 7 samples can be plotted per diagram. Diagrams created with ggvenn from the gglot2 R package.

**Supplemental Figure 3:**
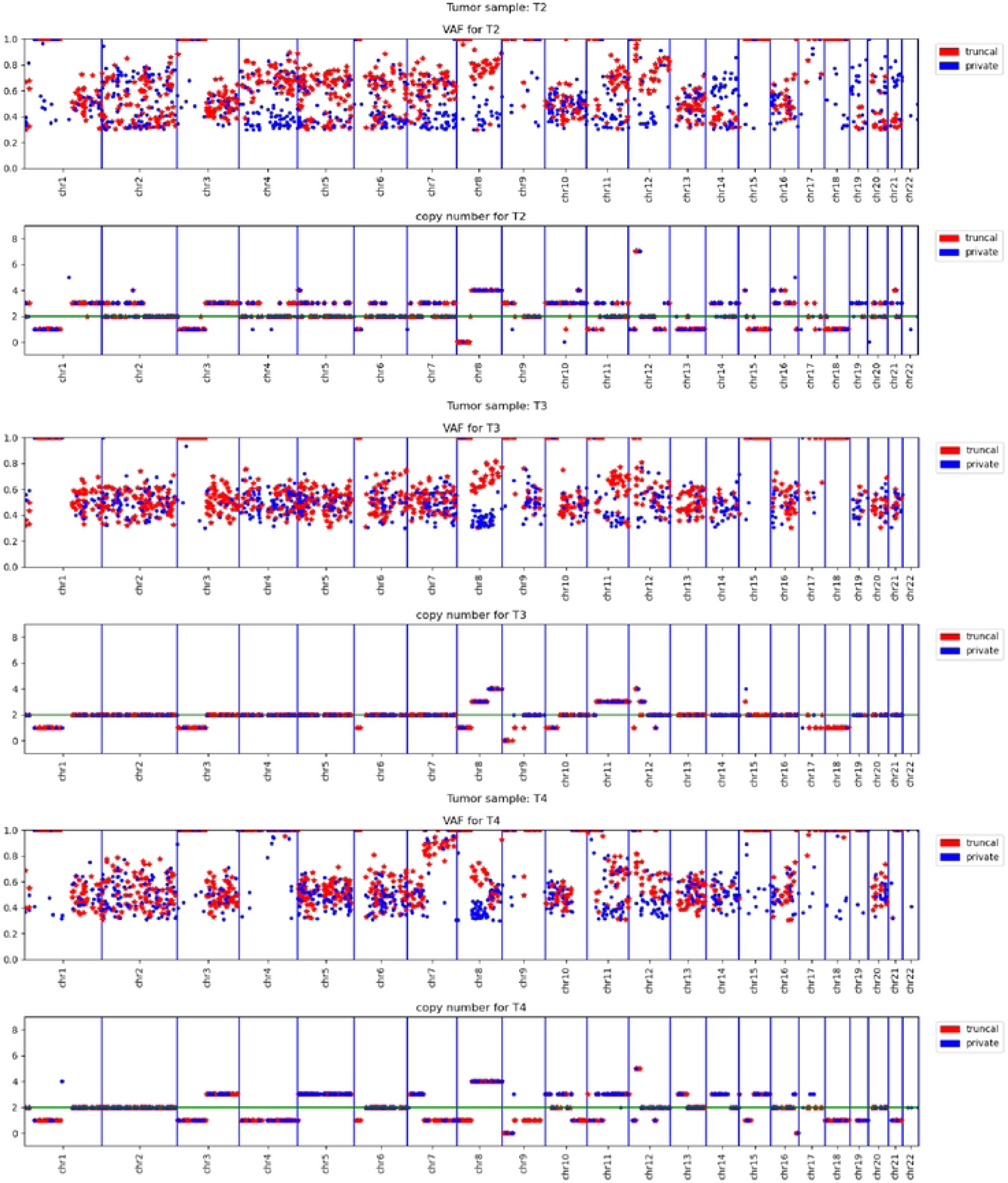
VAF and CN for truncal and private PAMs for WGS of A38 cell lines T2-4.

**Supplemental Figure 4:**
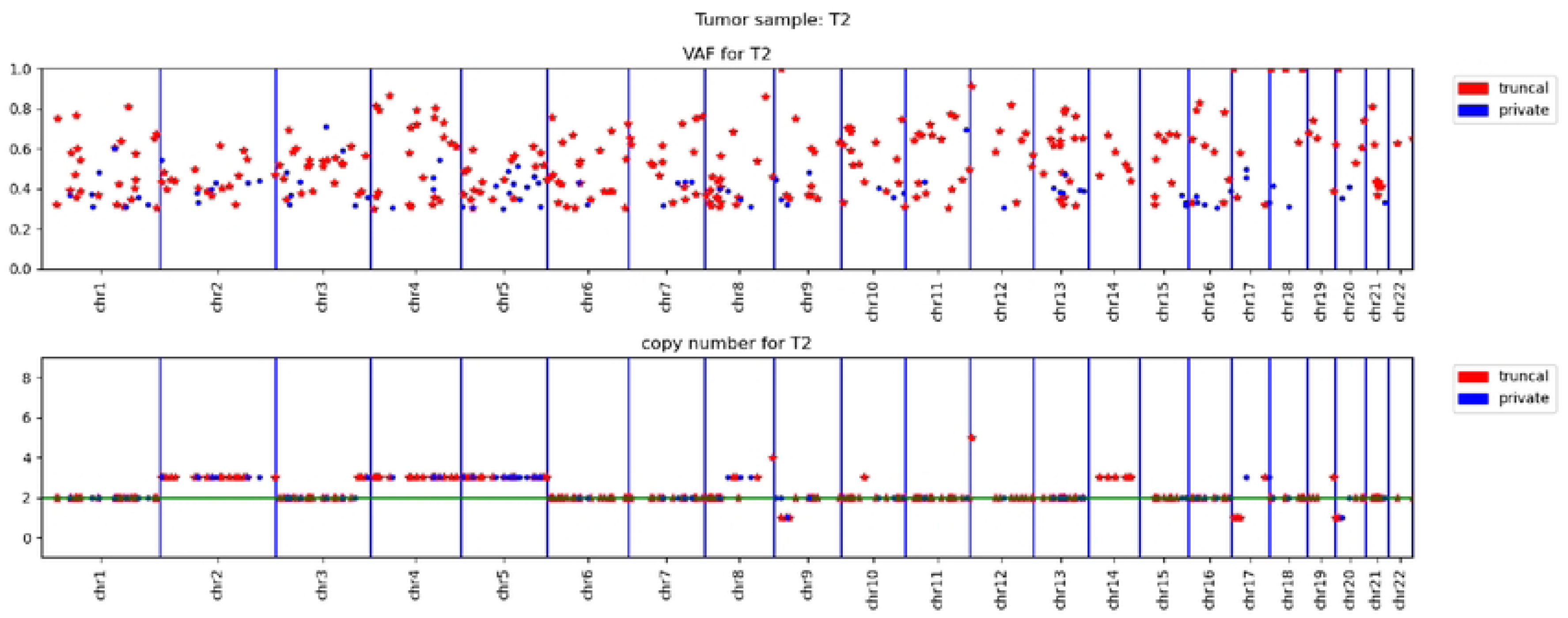
VAF and CN for truncal and private PAMs for WGS of A32 cell line T2.

**Supplemental Figure 5:**
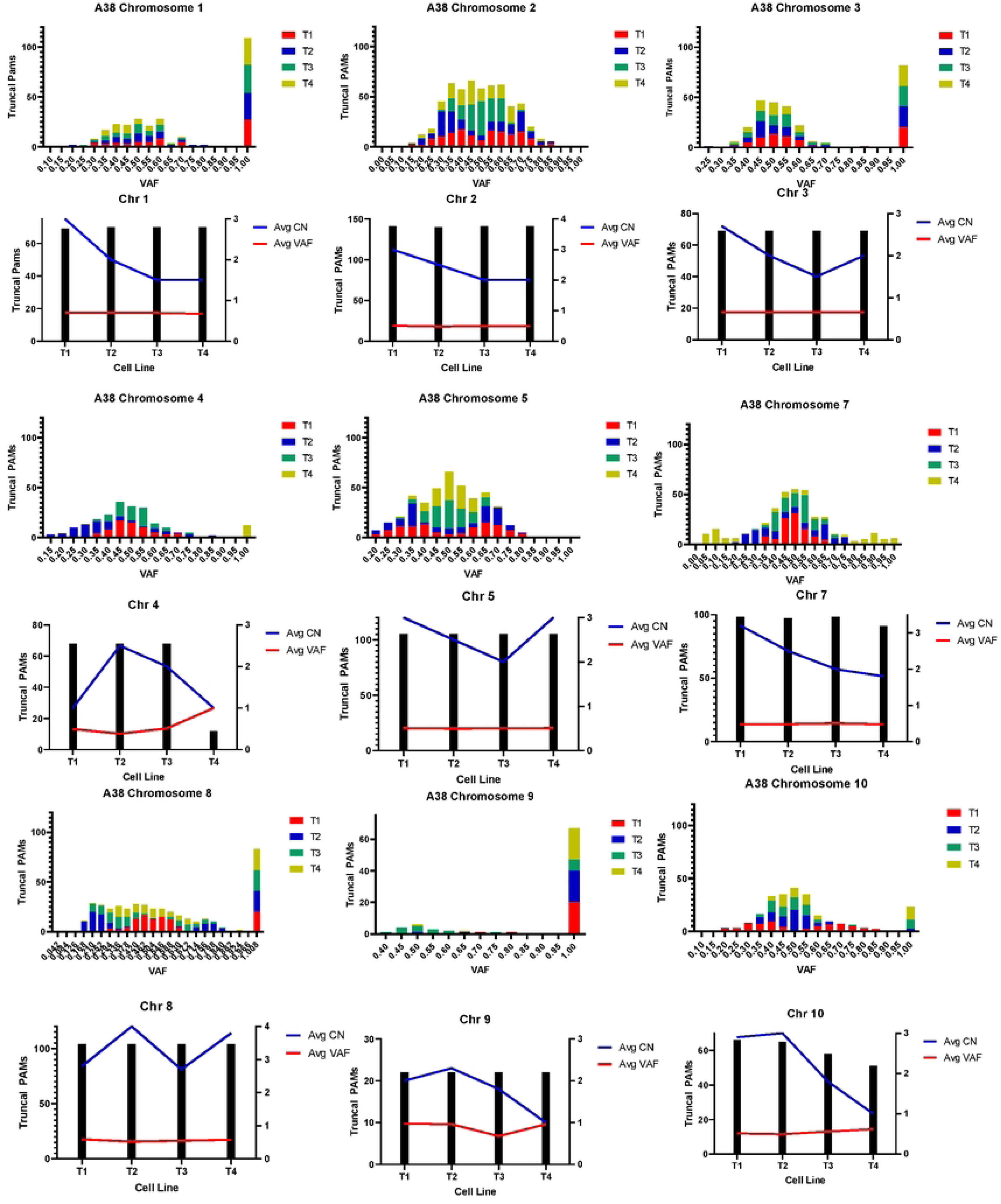
Histograms and CN/VAF analysis for truncal PAMs in A38 WGS cell line data, by chromosome.

**Supplemental Figure 6:**
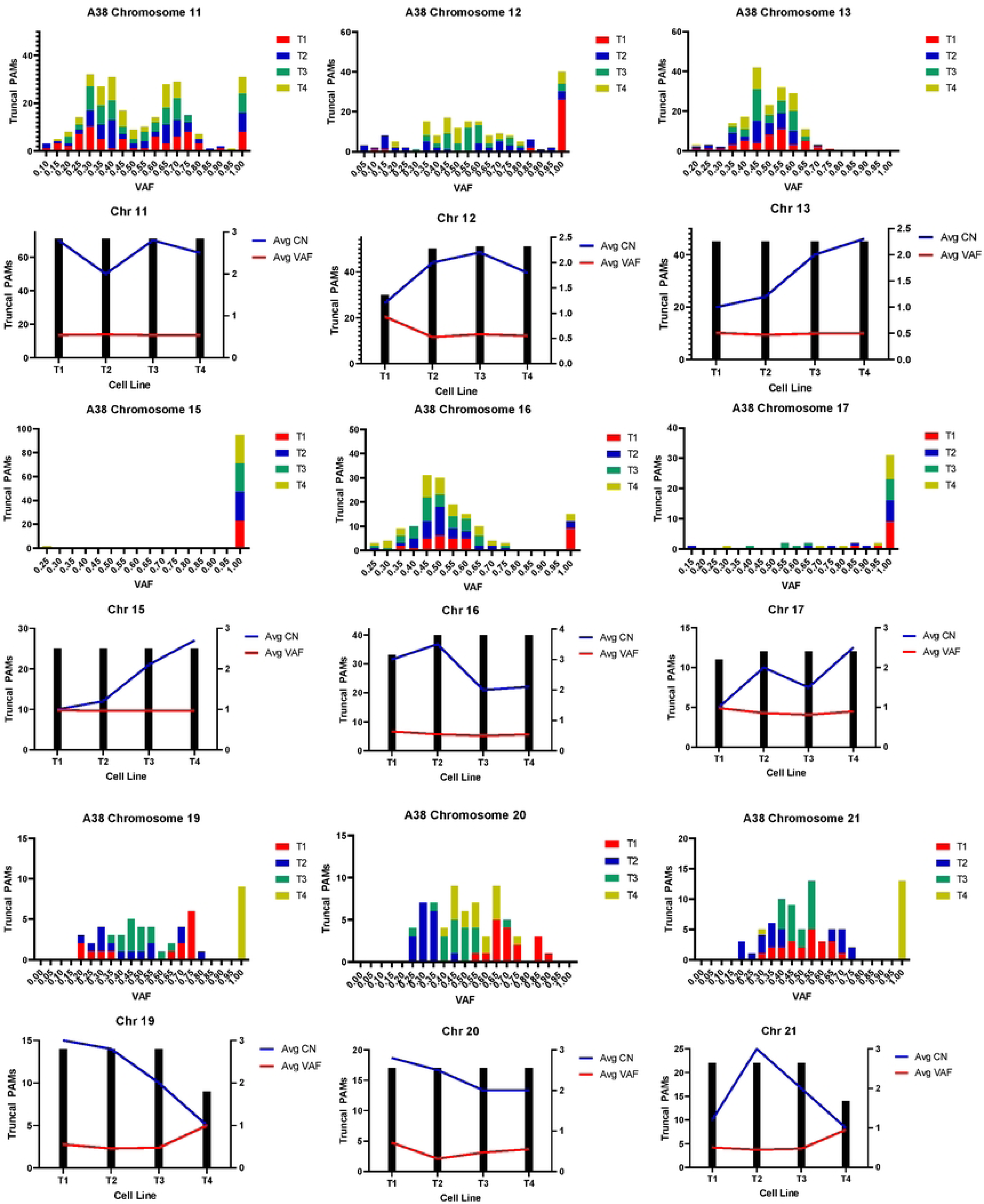
Histograms and CN/VAF analysis for truncal PAMs in A38 WGS cell line data, by chromosome.

**Supplemental Figure 7:**
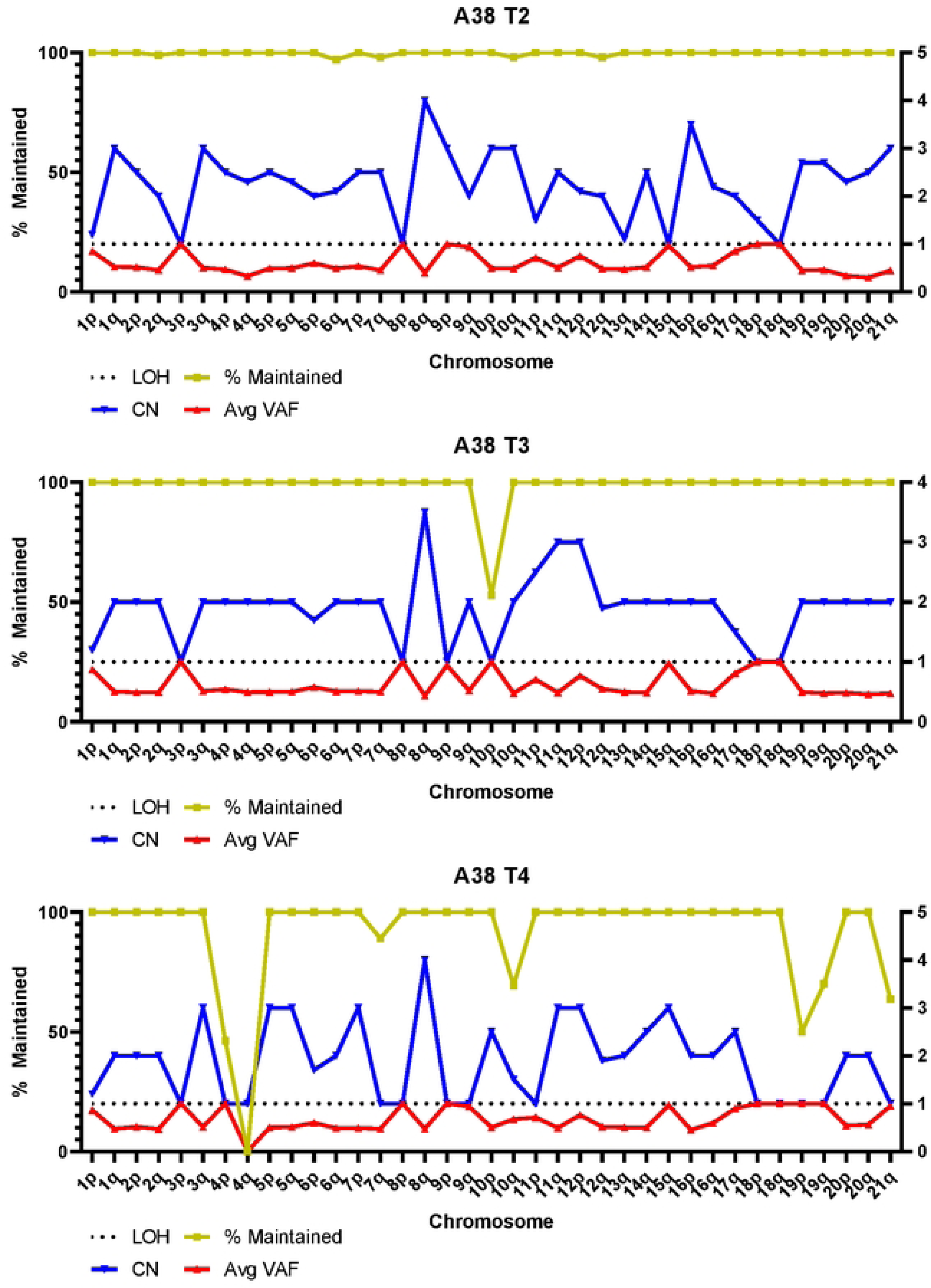
Plots of PAM maintenance, VAF, and CN for A38 cell lines T2-T4 analyzed for each chromosome arm.

**Supplemental Figure 8:**
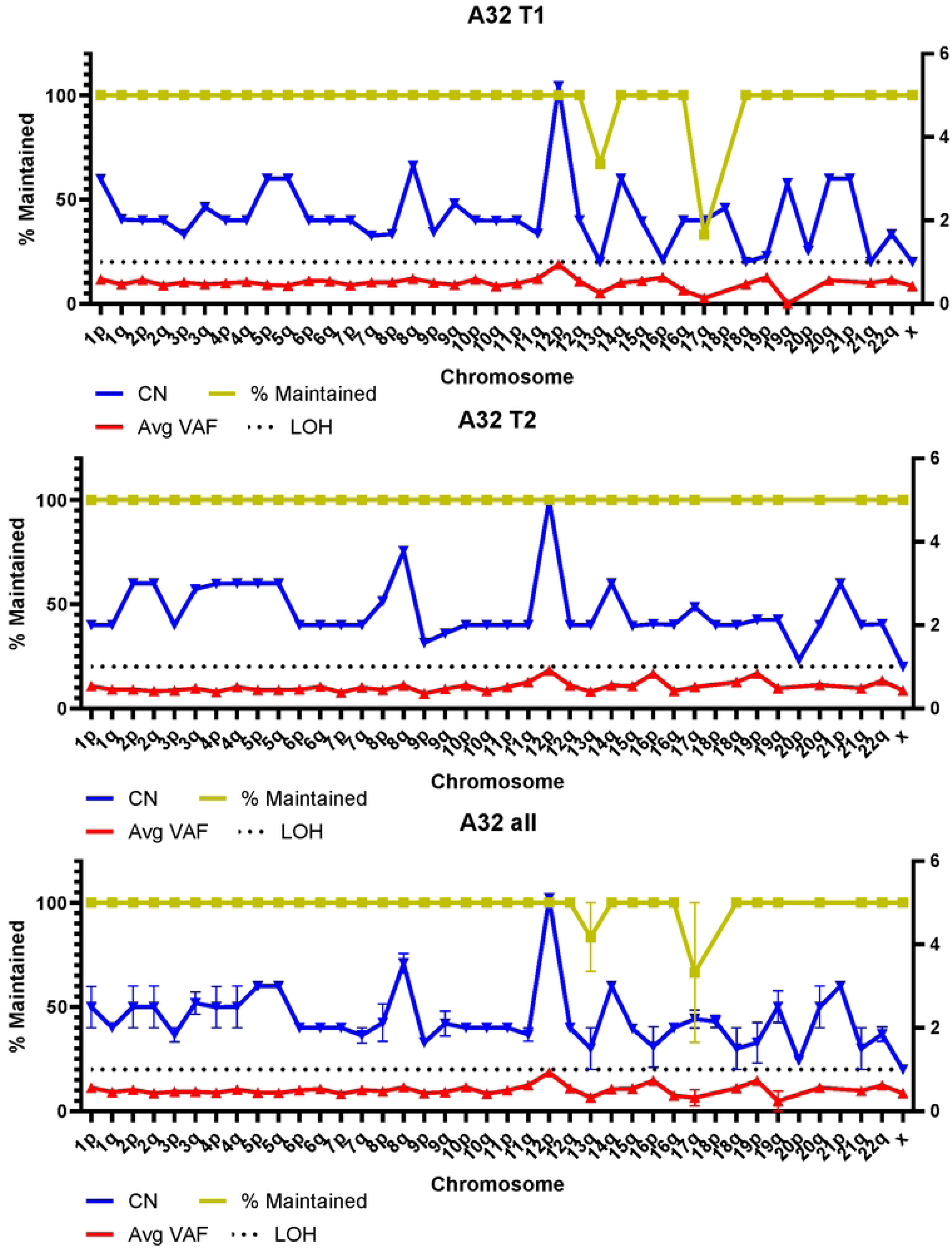
Plots of PAM maintenance, VAF, and CN for A38 cell lines T2-T4 analyzed for each chromosome arm.

**Supplemental Figure 9:**
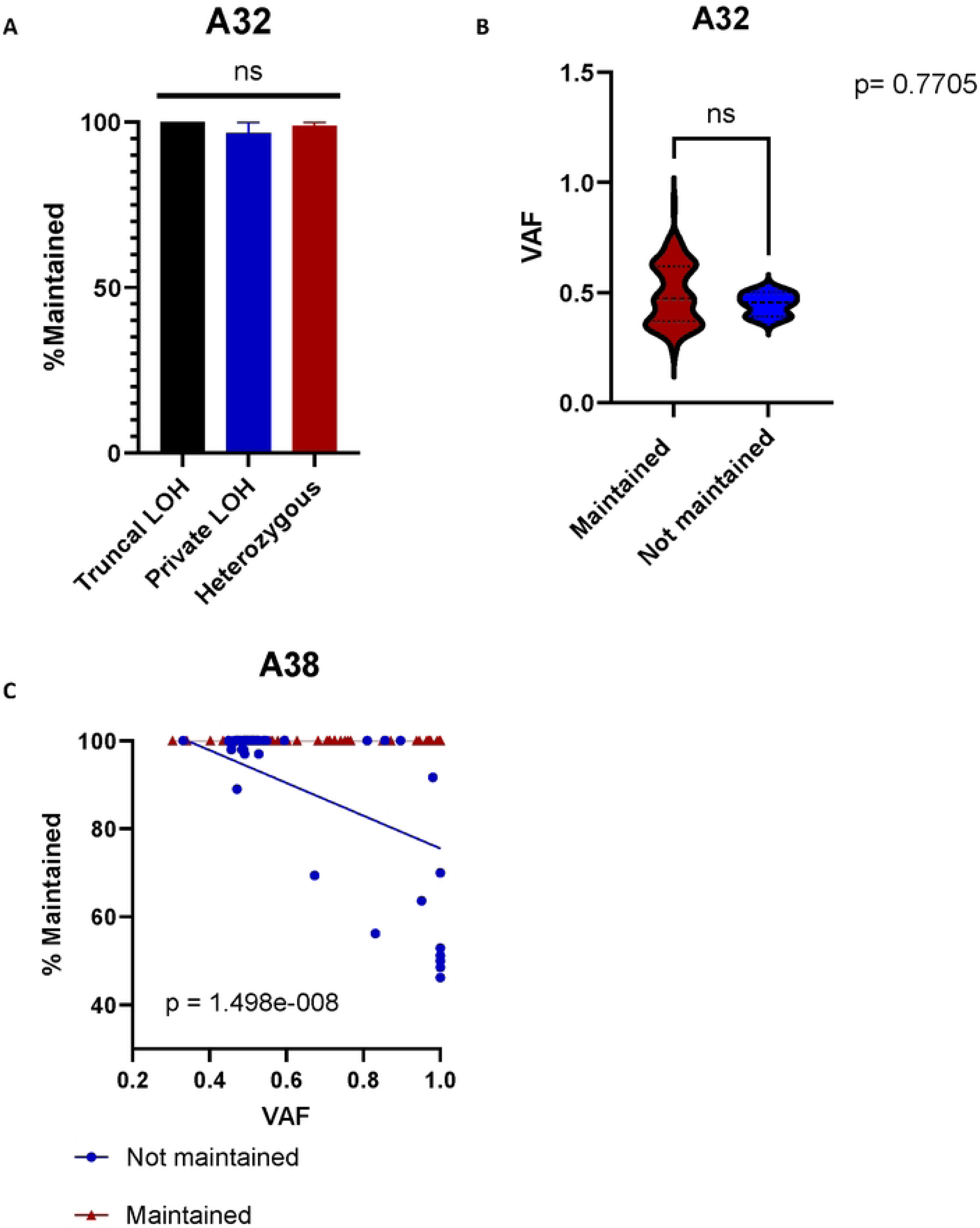
WGS was available for 2 cell lines and normal for case A32. A) No significant difference was found between the two cell lines for LOH and PAM maintenance (Kruskal- Wallis, p=0.3307) or for B) VAF between truncal PAMs maintained in both lines or lost in some (Mann-Whitney rank sum, p = 0.7705). n=165 for maintained PAMs, n=3 for PAMs lost in one cell line but not the other. C) Plot of truncal PAM VAF comparing PAMs maintained in all four cell lines (blue) and truncal PAMs lost in at least 1 cell line (red). A wide VAF distribution in present in PAMs present in all. For PAMs lost in some, VAFs cluster at 0.5 in samples where they are maintained, 1 when they are lost at a rate of 50%.

